# TgCep250 is dynamically processed through the division cycle and essential for structural integrity of the *Toxoplasma* centrosome

**DOI:** 10.1101/433466

**Authors:** Chun-Ti Chen, Marc-Jan Gubbels

## Abstract

The *Toxoplasma* centrosome is a unique bipartite structure comprising an inner- and outer-core responsible for the nuclear cycle (mitosis) and budding cycles (cytokinesis), respectively. These two cores remain associated during the cell cycle but have been proposed to function independently. Here, we describe the function of a large coiled-coil protein, TgCep250, in connecting the two centrosomal cores and promoting their structural integrity. Throughout the cell cycle TgCep250 localizes to the centrosome inner-core but resides on both inner- and outer-cores during the onset of cell division. This dynamic localization pattern is associated with proteolysis: the processed version residing on the inner-core. In the absence of TgCep250, stray centrosome inner- and outer-core foci were observed; detachment of the inner-outer-core connection resulted in nuclear partitioning defects. The detachment between centrosome inner- and outer-core was found in only one of the centrosomes during cell division, indicating distinct states of mother and daughter centrosomes. We further dissected the hierarchical organization of centrosome and kinetochore complex through depletion of kinetochore component TgNuf2, which resulted in dissociation of the intact bipolar centrosome from the nuclear periphery. Together, these data suggest that TgCep250 bridges the interaction between the centrosome cores but not between the inner-core and kinetochore.

**Short Summary:** The opportunistic apicomplexan parasite *Toxoplasma gondii* uses a bipartite centrosome to independently regulate mitosis and cytokinesis. Here we report a large coiled-coil protein that functions to integrate the two centrosomal cores for faithful cell division. This study also reveals the layered structural organization of the centrosome/kinetochore complex.

## Introduction

The phylum Apicomplexa consists of almost exclusively obligate intracellular parasites, which have significant impact on public health. Among these single-celled organisms, *Toxoplasma gondii* is one of the most successful zoonotic pathogens that can be transmitted and replicate in many different host species. *T. gondii* has developed flexible replication strategies allowing efficient survival in distinct tissue types. Tachyzoites are the acute replication form of *T. gondii* and are responsible for most of the clinical burden in infected individuals. They undergo a binary replication mode termed endodyogeny where two daughter cells assemble internally and emerge from the mother cell. The cell cycle follows the sequence of G1, S and M phases, but the G2 phase is absent or too short to be detected ^1^ and is driven by a set of unconventional CDK-related kinases (Crks) and cyclins ^2^. Mitosis is (semi-)closed and tightly spatially and temporally coordinated with cytokinesis as these two events occur concomitantly. The 14 chromosomes are clustered at the centromeric region throughout the cell cycle at the ‘centrocone’, a unique membranous structure that houses spindle microtubule during mitosis ^3^. The tight organization of chromosomes is thought to ascertain each daughter cell receives a full set of genetic material when the parasite undergoes more complex division cycle in other cell cycle stages ^4^. In its definitive feline host, endopolygeny consists of several rounds of DNA synthesis and mitosis in the same cytoplasmic mass followed by synchronized cytokinesis in accordance with the last round of mitosis, producing 8-16 uni-nucleated progeny per replication cycle ^5^.

The *Toxoplasma* centrosome is divergent from mammalian cells in architecture and composition. For example, the centrioles are composed of nine singlet microtubules, smaller in size than mammalian centrioles, and the centriole pair is arranged in parallel rather than perpendicular ^4^. Orthologs of many key components in the mammalian centrosome cannot be found in the *Toxoplasma* genome ^6^. Even without canonical protein orthologs, recent studies have shown that a group of coiled-coil proteins localize to the centrosome and support proper karyokinesis and cytokinesis ^7–8^. This work identified a distinctive architecture of two centrosome cores: the outer-core (distal to the nucleus) and the inner-core (proximal to the nucleus).

The *Toxoplasma* centrosome is the key organelle coordinating mitotic and cytokinetic rounds. The centrosome resides at the apical end of the nucleus during interphase and remains closely associated with a specialized nuclear envelope fold termed the centrocone that houses the spindle microtubules (MTs) during mitosis ^9^. The centrosome also plays a critical role in partitioning the single copy Golgi apparatus and the chloroplast-like organelle, the apicoplast, during cell division ^10^. Late in G1, the centrosome rotates to the distal end of the nucleus where it duplicates and then returns to the proximal end ^11^. At this point, the spindle MTs assemble and are stabilized within the centrocone, whereas subsequently the centrosome serves as a scaffold for assembly of daughter cell cytoskeletal components ^12^. In mitosis, a striated fiber assemblin (SFA) protein emanates from the centrosome and connects the daughter apical conoid to the centrocone. SFA polymerization is thought to maintain the stoichiometric count of daughter cells and replicating genomic materials. In SFA depleted cells, cell division is completely disrupted ^13^. Throughout these timely regulated events, little is known about centrosome biogenesis or regulation in *Toxoplasma.* Although orthologs in charge of new centriole biogenesis, Polo-like kinase PLK4 and its substrate STIL/Ana2/SAS5, are absent from the *Toxoplasma* genome, previous work has shown that a conserved NIMA-kinase, TgNek1, is required for centrosome splitting ^14^ and a TgMAPK-L1 place a role in inhibiting centrosome over-duplication ^7^. However, exactly on what aspects of the centrosome these kinases act and what their substrates are remains unclear.

Here we asked whether TgCep250 could be the substrate of TgNek1 required for centrosome splitting in orthogology to the human HsNek2 - C-NAP1/Cep250 relationship in centrosomes splitting ^15^. Our data reject this hypothesis but instead revealed that TgCep250 is required to connect the inner-core with the outer-core as well as to maintain overall centrosome integrity. Moreover, our data are the first report directly hinting at a differential status of mother versus daughter centrosome in the Apicomplexa. In conclusion, TgCep250 is not the critical TgNek1 substrate, but its function in keeping centrosome complexes connected appears to be orthologous to the role of human C-NAP1/Cep250.

## Results

### TgCep250 has a dynamic localization and is associated with proteolysis

To start charting the function of previously reported *Toxoplasma* centrosome protein TgCep250 (TGGT1_212880) ^7^ we first characterized the subcellular localization of TgCep250 throughout the parasite’s tachyzoite division cycle. TgCep250 was tagged endogenously with a C-terminal YFP tag ^16^ and the TgCep250-YFP parasite line was transfected with a fosmid expressing TgCep250L1-HA as an inner-core centrosome marker ^7^. During the G1 phase of the cell cycle TgCep250-YFP is closely associated with the centrosome inner-core (highlighted by TgCep250L1) whereas during cytokinesis we observed four TgCep250-YFP foci where the upper two foci in close apposition to the outer-core (Fig. 1).

**Figure 1.**
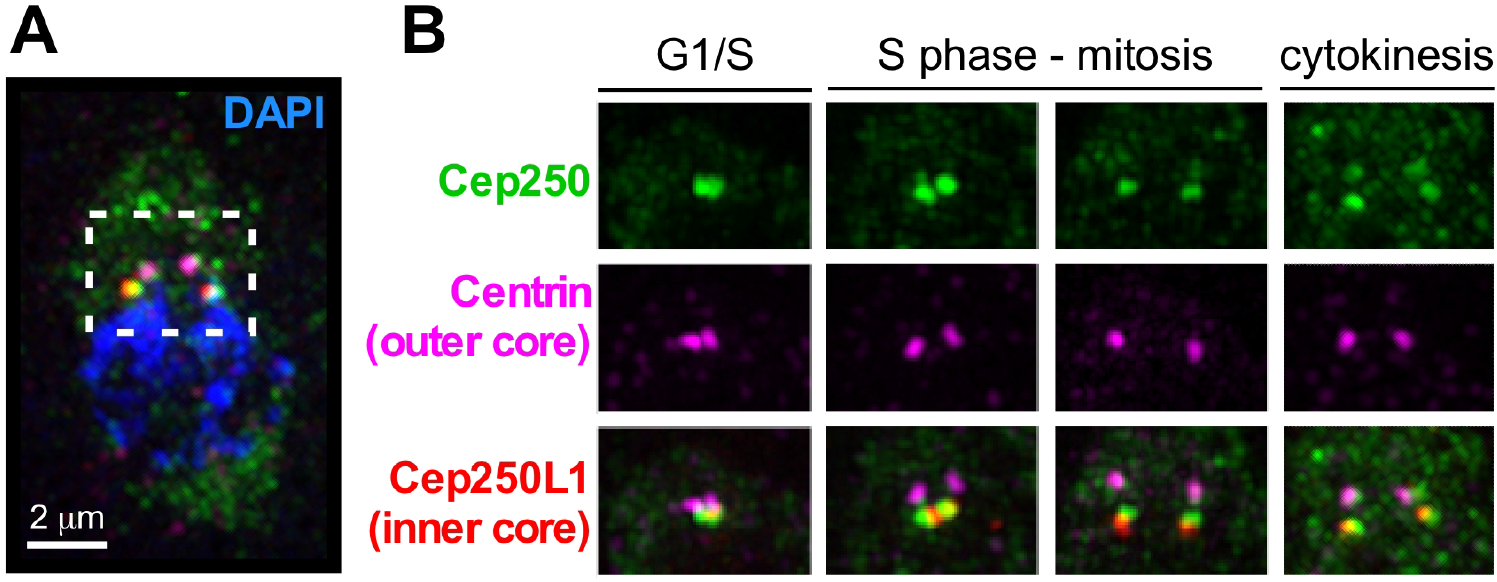
Dynamic localization of TgCep250 throughout the cell cycle. A) IFA of TgCep250-YFP parasites co-stained with α-GFP (Cep250, shown in green), α-Centrin (centrosome outer-core marker, shown in magenta), α-HA (centrosome inner-core marker shown in red) and DAPI (in blue). Images acquired by super-resolution microscopy (SIM). B) Single-colored panels showed dynamic localization of TgCep250-YFP during asexual division cycle. Cell cycle stages as indicated.

To dissect the function of TgCep250, we generated a conditional knockdown line (TgCep250-cKD) using a tetracycline regulatable promoter. Simultaneously, we inserted a single-Myc tag at the N-terminus of the gene (Fig. 2A). Correct integration of the construct through single homologous recombination was validated by PCR (Fig. 2B). In the presence of anhydrotetracycline (ATc), Myc-TgCep250 protein becomes undetectable within six hours of incubation by immunofluorescence assay (IFA) and western blot (Fig. 2C and D). Interestingly, the N-terminal Myc-tagged TgCep250 only formed two foci co-localizing with α-Centrin throughout the cell cycle. This suggested that the placement of the tag on TgCep250 either alters the subcellular localization or that proteolytic processing is associated with this differential localization pattern. To test this hypothesis, we added a C-terminal 3xTy tag to the Myc-TgCep250-cKD locus (Fig. 3A) and analyzed TgCep250-3xTy by IFA and western blot. As shown in Figure 3B and 3C, C-terminally 3xTy tagged TgCep250 forms four foci like TgCep250-YFP. Moreover, we observed multiple protein bands smaller than the full-length protein on the blot using α-Ty antibody (Fig. 3C; note that full-length TgCep250 is a large coil-coiled protein with a molecular weight predicted to be 762 kDa). Thus, these data support the notion that that only full-length TgCep250 localizes to the outer-core (during the onset of cytokinesis) and a truncated version of TgCep250 localizes to the inner-core (throughout all cell cycle stages; Fig. 3B and 3D). In line with this hypothesis, we only detected a single band on western blot consistent with the full-length protein using α-Myc antibody (Fig. 2D). Thus, we conclude that TgCep250 has dynamic localization pattern during the cell cycle and that protein processing appears to affect association of TgCep250 to the centrosome inner cores.

**Figure 2.**
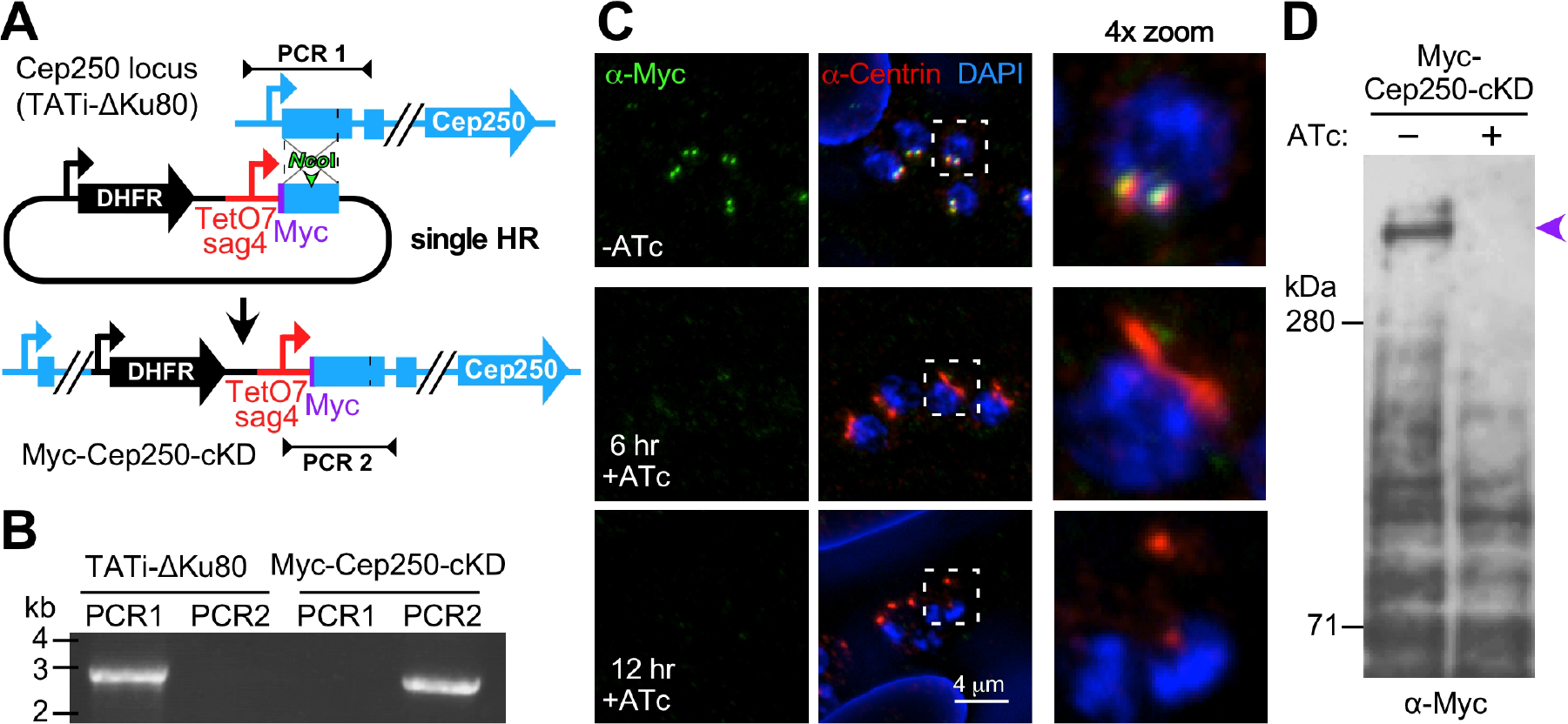
Generation and validation of a TgCep250-cKD line. A) Schematic representation of TgCep250 conditional knockdown (cKD) construction by single homologous recombination. B) PCR validation using TATi-ΔKu80 (parent) and Myc-TgCep250-cKD genomic DNA as templates. Primer pairs as indicated in Fig. 2A. C) IFA using α-Myc and α-Centrin showed that the Myc-tagged TgCep250 is depleted with in 6-hour incubation of ATc. Boxed areas magnified in the 4× zoom panels. D) Western blot using α-Myc showed that the protein expression level is down-regulated in the presence of ATc. The purple arrowhead marks the full-length Cep250 protein (note that TgCep250 has a predicted molecular weight of 762 kDa); the smaller bands present in both the − and +ATc samples and considered aspecific and serve as loading control.

**Figure 3.**
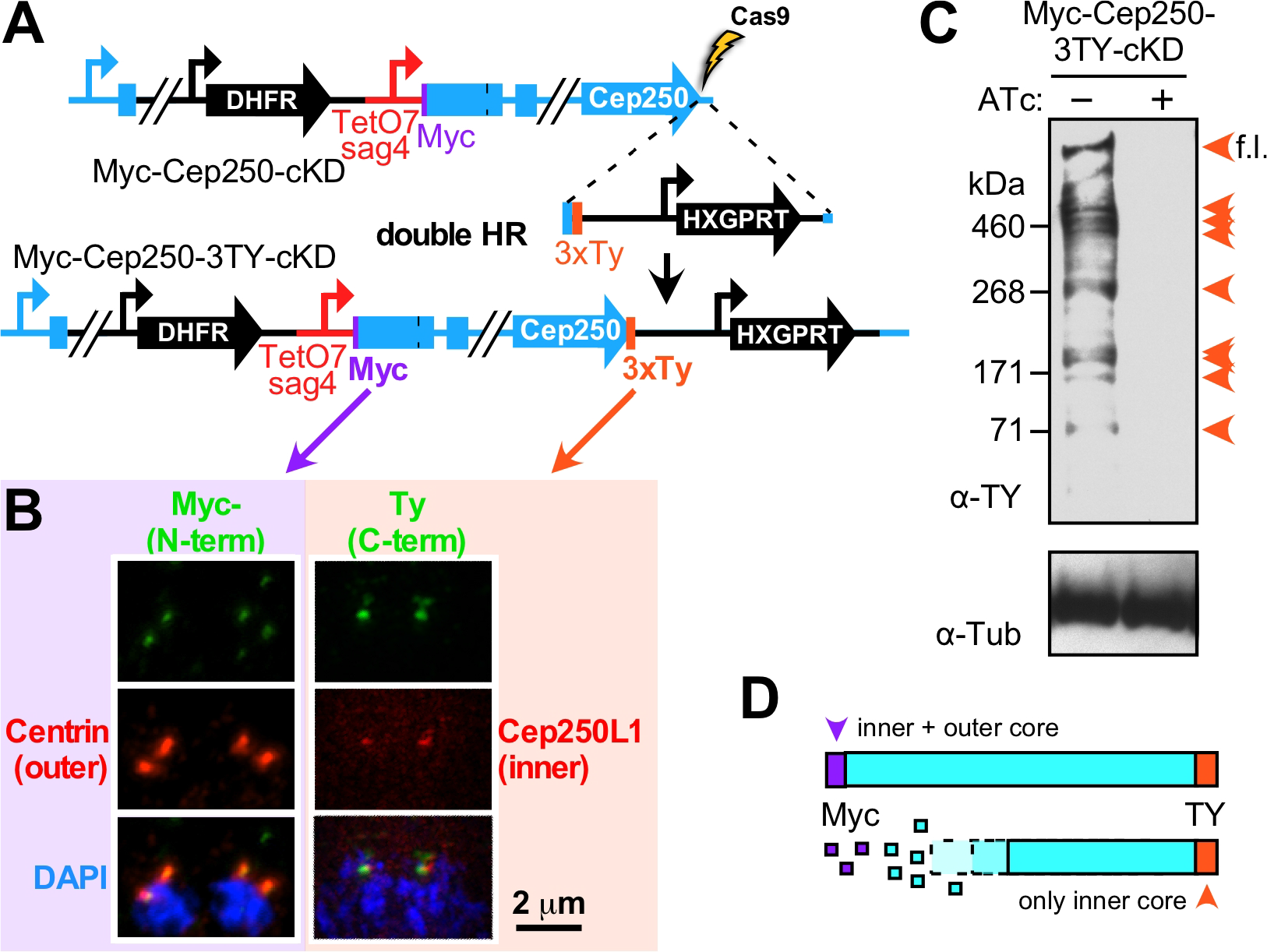
Differential localization of TgCep250 is regulated by proteolysis. A) Schematic representation of Myc-TgCep250-3xTy-cKD construction using CRISPR/Cas9. B) Myc-TgCep250-cKD localized only to the outer-core (stained with α-Myc and α-Centrin) and TgCep250-3xTy localized to both inner- and outer-core (stained with α-Ty and α-HA on endogenously-tagged TgCep250L1-HA as inner-core marker). Left panels are conventional wide-field microscopy; right panels are super-resolution (SIM) microscopy. C) Multiple protein bands were detected in western blot using α-Ty antibody suggestive of extensive proteolytic processing. α-tubulin antiserum (12G10) was used as loading control. Orange arrowheads mark the specifically detected TgCep250 protein fragments. D) Schematic interpretation of western blot and IFA data representing the relationship between TgCep250 proteolysis and subcellular localization.

### TgCep250 is essential and links the centrosome inner- and outer-cores

Next, we probed the function of TgCep250 with a series of experiments using the TgCep250-cKD parasite line. In the presence of ATc, TgCep250-cKD parasites failed to form plaques suggesting TgCep250 is essential for parasite survival (Fig. 4A). Phenotypic analysis by IFA showed that in absence of TgCep250 the nucleus failed to partition to the daughter cells (Fig. 4B, white arrow) and as a result anuclear parasites were observed (Fig. 4B, white arrowheads). The stoichiometric balance of 1:1 daughter to (outer-)centrosome count was also disrupted resulting in three Centrin foci accumulated in a single cell boundary highlighted by IMC3 (Fig. 4B, double arrowhead). To examine the nuclear partitioning defect in more detail, we used different sets of antibodies to analyze TgCep250 depleted parasites. As shown in Figure 4C, we observed and quantified three different centrosomal defects in TgCep250 depleted cells using centrosome inner-(TgCep250L1) and outer-core (Centrin) markers. We observed detachment of inner-core from the outer-core in the majority of mitotic, mutant parasites (marked in yellow). However, we also observed parasites wherein the outer-core split into two foci while only two inner-cores were present, suggestive of undergoing an additional replication cycle (marked in cyan; 10% of the mitotic parasites). Finally, about 35% of the mitotic parasites displayed, stray inner core foci (marked in magenta) which is a combination of the two other phenotypes. Detachment of inner- and outer-cores likely results in the nuclear partitioning defect observed in Figure 4B. An additional observation is that detachment of inner- and outer-cores occurred in most instances on only one of the two centrosomes. The most logical explanation is that the detached inner/outer-core pair represents the daughter centrosome and the intact pair represents the mother centrosome ^17^. In this scenario, during centrosome biogenesis, TgCep250 cannot be incorporated into the newly formed daughter centrosome resulting in a detached inner- and outer-core. However, the TgCep250 in the mother centrosome is stably incorporated thereby maintaining inner- and outer-core adhesion in this centrosome.

**Figure 4.**
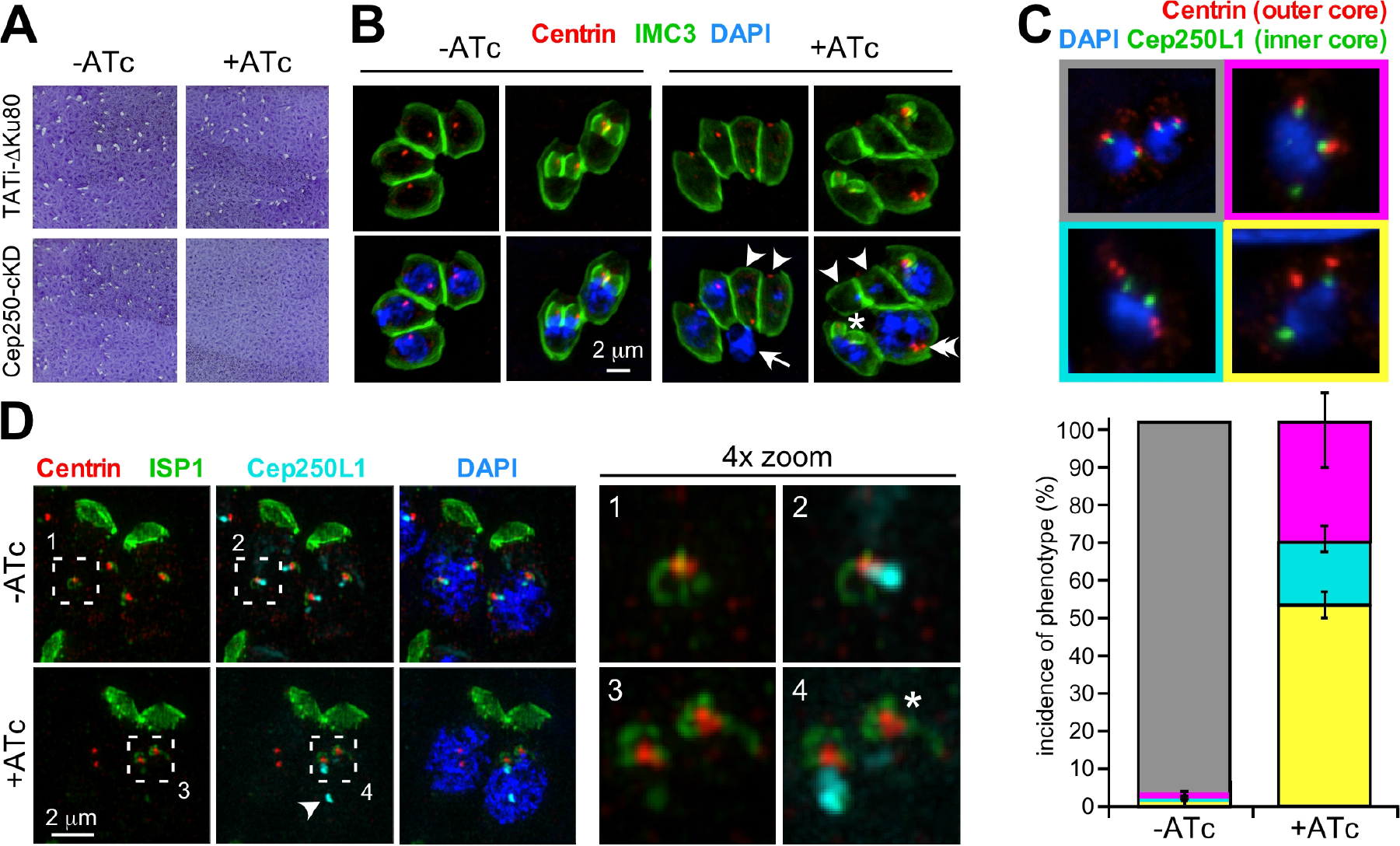
Loss of TgCep250 causes lethal centrosomal defects. A) Plaque assay of TgCep250-cKD and the TATi-ΔKu80 parent line ±ATc for seven days shows that TgCep250 is essential. B) IFA of TgCep250-cKD stained with α-Centrin (in red) and α-IMC3 (in green) ±ATc. Arrowheads mark parasites with nuclear loss; arrow marks a nucleus outside a parasite; double arrowhead marks a parasite with two nuclei and 3 outer-cores.; asterisk marks a parasite with daughter buds out of sync with the rest of the vacuole. C) TgCep250-cKD parasites co-expressing TgCep250L1-HA were stained with α-Centrin (in red) and α-HA (in green) marking the outer- and inner-cores, respectively. Top panel shows IFA images representative of observed phenotypes: grey boxed: control wild-type nuclei displaying duplicated centrosomes; cyan boxed: outer-core duplication independent of inner-core duplication; magenta boxed: stray inner-core complex in presence of two normal appearing inner/outer pairs; yellow boxed: complete detachment of inner- and outer-cores. Lower panel: quantification of defective centrosome phenotypes; bar colors match the colored boxes in the top panel. At least 100 nuclei with a duplicated centrosome pair were counted; error bars represent standard deviation from 3 independent experiments D) Super-resolution (SIM) IFA of TgCep250-cKD ±ATc co-expressing TgCep250L1-HA stained with α-ISP1 (green: early daughter scaffold marker), α-Centrin (red: outer-core) and α-HA (cyan: inner-core). Numbered, boxed areas are magnified on the right. Note that outer-cores are normally associated with daughter scaffolds (asterisk), even if inner-core connection is lost (arrowhead).

Interestingly, the ability to form daughter buds remained intact in TgCep250 depleted cells (Fig. 4B; daughter buds marked with asterisk). Thus, we sought to determine whether the detached centrosome cores remain functional in daughter bud assembly. For this purpose, an IMC-subcompartment protein ISP1 was used as a marker to determine early daughter formation near the centrosome ^18^. As shown in Figure 4D, α-ISP1 signal was detected surrounding the outer-core in the presence and absence of ATc, even when the outer-core is detached from the inner-core (Fig. 4D 4× zoom, asterisk). These results indicate that the function of the outer-core in daughter bud assembly remains intact when it is detached from the inner-core. This observation is in line with the proposed working model predicting that the inner-core and outer-core functions can act independently ^7, 19^.

### TgCep250 is not required for the connection of the inner-core with the kinetochore

Since the anuclear phenotype observed in TgCep250-cKD parasites was very similar to the anuclear daughters observed in parasites depleted in kinetochore component TgNuf2 ^20^ we tested how these two observations relate to each other. We performed IFA assays using mitotic markers to reveal the interactions between centrosome outer-core (α-Centrin), the kinetochore (α-TgNdc80) ^20^ and the spindle microtubules (EB1-YFP) ^11^. As shown in Figure 5, in the absence of TgCep250, both kinetochore complex and spindle microtubules became detached from the centrosome outer-core (Fig. 5A, white arrowhead). To determine whether TgCep250 plays a role in interaction between the inner-core and the kinetochore we co-stained the parasite with inner-core (TgCep250L1) and kinetochore (TgNdc80) markers. Surprisingly, although TgCep250L1 fragmented into multiple foci, the majority of TgNdc80 foci co-localized with TgCep250L1 foci in the absence of TgCep250, even though in some cases the connection with the nucleus was gone (Fig. 5B). Thus, the inner-core and the kinetochore remained associated in the absence of TgCep250 and indicates that the nuclear function of the inner core is also maintained when the inner/outer-core association is lost.

**Figure 5.**
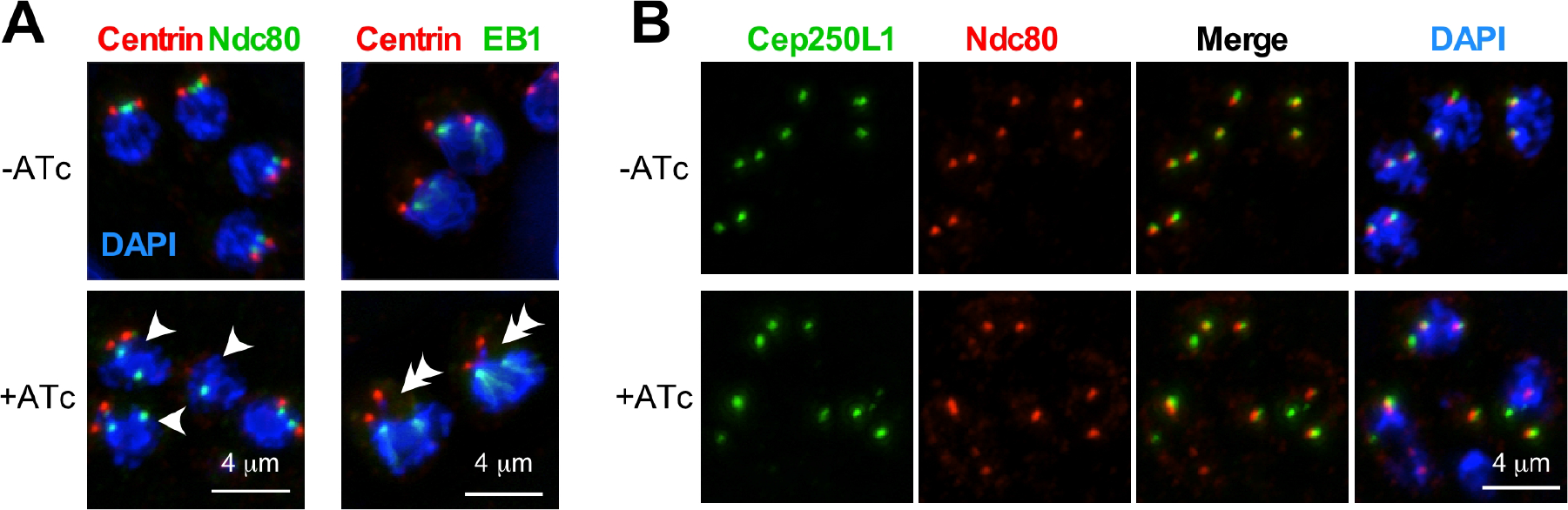
The inner-core remains associated with the mitotic apparatus in TgCep250 depleted parasites. A) IFA of TgCep250-cKD ±ATc stained with α-Centrin (outer-core; red) and co-stained with either α-TgNdc80 (kinetochore; green; left panels) or co-expression of TgEB1-YFP (spindle microtubules; green; right panels). Arrowheads mark nuclei displaying loss of outer-core association with the kinetochore; double arrowheads mark nuclei displaying loss of outer-core association with the spindle. B) IFA of TgCep250-cKD ±ATc expressing TgCep250L1-HA (inner-core: α-HA; green) and co-stained α-TgNdc80 (kinetochore; red) showed that the interaction between the kinetochore and inner-core centrosome remains intact in the absence of TgCep250.

### TgNuf2 anchors the centrosome to the nuclear periphery

To further dissect the association between the inner-core and the kinetochore, we expressed the inner-core marker TgCep250L1 in a kinetochore-depleted parasite line, TgNuf2-cKD ^20^. It has been reported that in the absence of TgNuf2, the association between the centrosome and spindle pole is abolished and the centrosome is pulled away from the nucleus ^20^. In Figure 6, we observed that the centrosome inner- and outer-core co-migrate away from the nucleus (Fig. 6 arrowhead). In this scenario, association between outer-core and inner-core remains intact in the absence of TgNuf2. These data suggest that the kinetochore is required to anchor the centrosome cores to the nuclear periphery but not required for the connection between the inner- and outer-cores.

**Figure 6.**
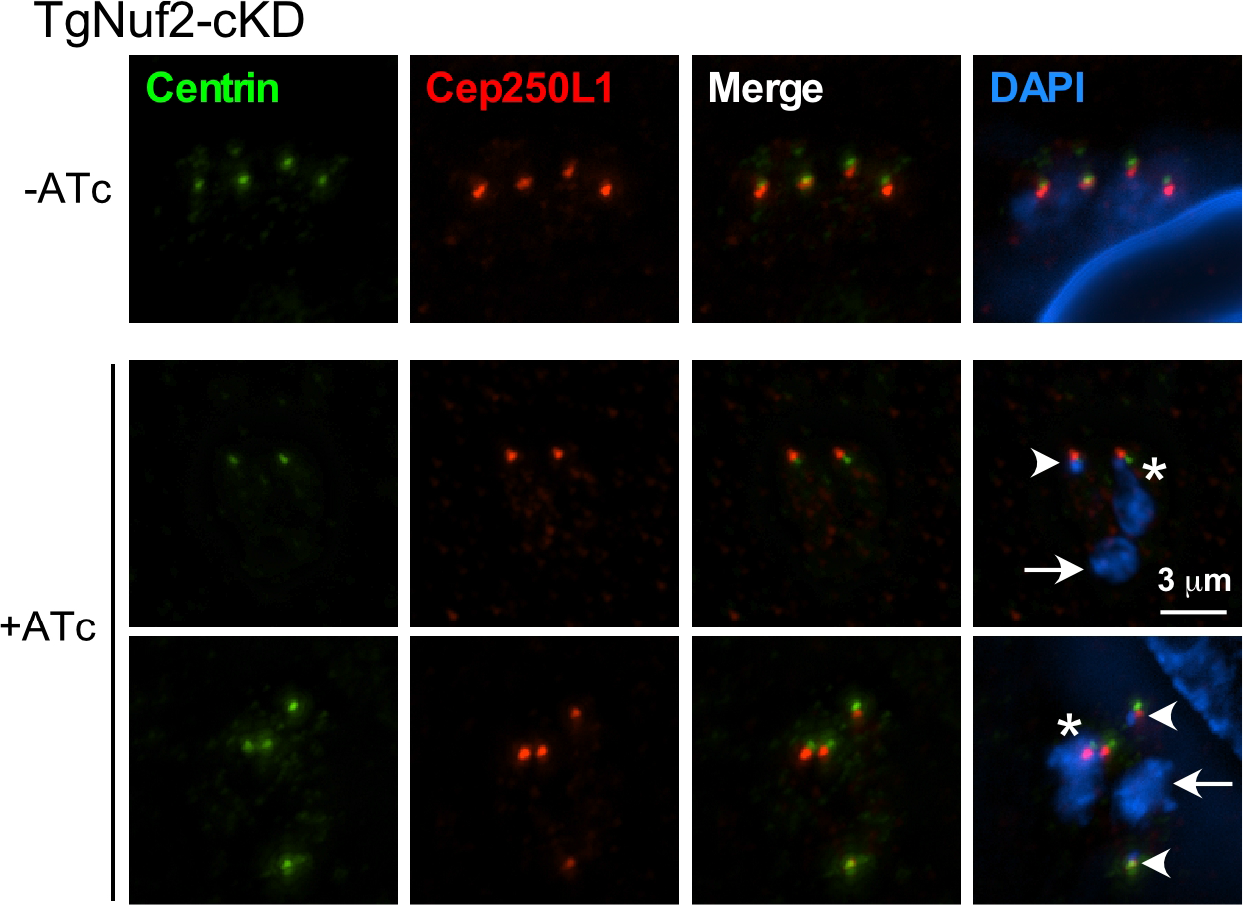
The kinetochore is required to anchor centrosome cores to the nuclear periphery. TgNuf2-cKD parasites treated ±ATc stained with centrosome outer-core (α-Centrin, in green) and inner-core (TgCep250L1-HA; α-HA, in red;) markers. Asterisks mark intact nucleus-centrosome connections; arrowheads mark intact inner/outer-core pairs strayed from the nucleus (the associated small DAPI spot is the plastid genome); arrows mark mis-segregated nuclei.

### TgNek1 promotes separation of only the outer-cores

It has been shown that a *Toxoplasma* ortholog of NIMA kinases, TgNek1, is responsible for centrosome separation in tachyzoites ^14^. To dissect the regulation of centrosome core separation in a TgNek1 depleted background we constructed a conditional knockdown line of TgNek1 (TgNek1-cKD) (Fig. S1A). Plaque assays showed that in the presence of ATc, TgNek1-cKD parasites display a severe growth defect (Fig. S1B) consistent with the temperature sensitive TgNek1 phenotype previously reported ^14^. Moreover, IFA with α-TgNek1 confirmed that TgNek1 expression is depleted in the presence of ATc (Fig. S1C). Interestingly, upon depletion of TgNek1 the duplicated outer-cores remain connected whereas the inner-cores separate along with the kinetochores (Fig. 7). The distance between separated kinetochores in TgNek1-cKD and wild type parasites is comparable, which in turn is comparable with previously reported distances ^7^. This indicates that the spindle MT polymerization is not affected when outer-core splitting is blocked. In all, we concluded that the centrosome outer-core, inner-core and kinetochore are hierarchically organized and that TgCep250 is responsible for connecting the inner- and outer-core, but not the inner-core and the kinetochore.

**Figure 7.**
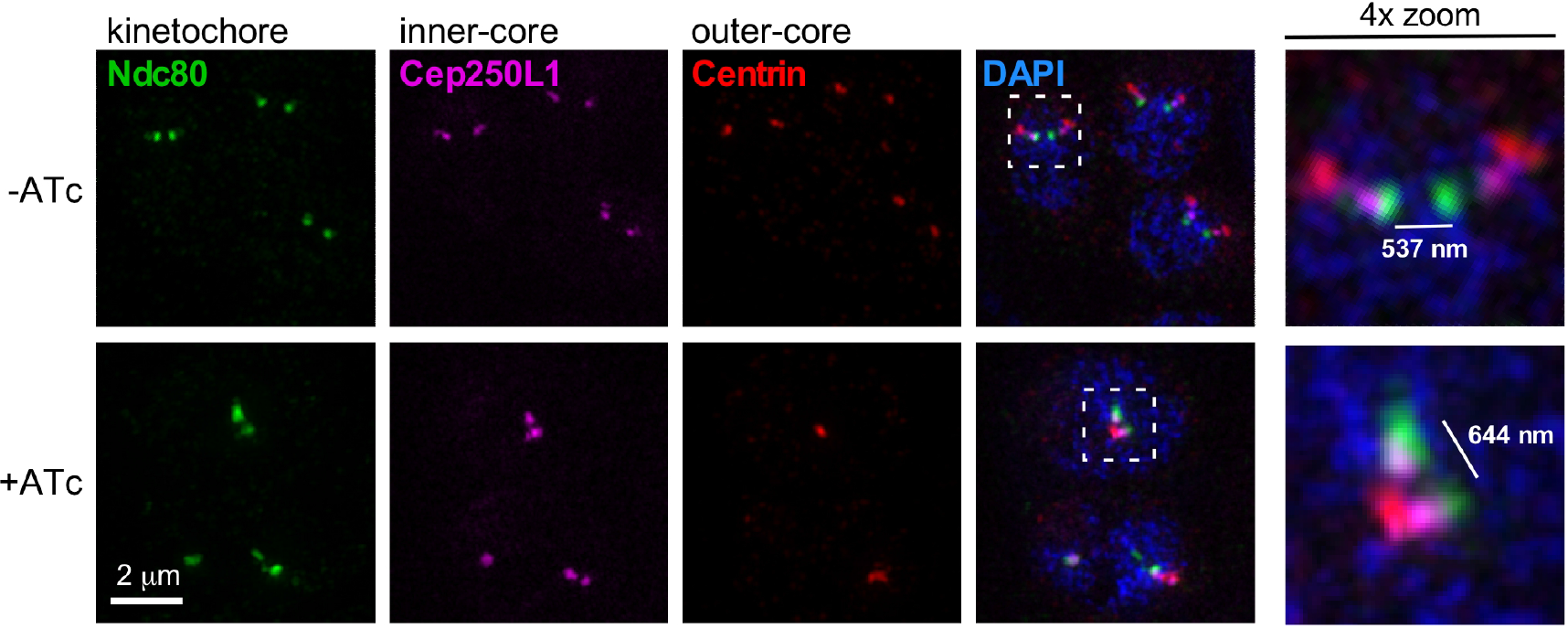
Loss of TgNek1 prevents outer-core splitting but does not interfere with inner-core separation. Image acquired by super-resolution microscopy (SIM) of TgNek1-cKD parasites ±ATc using kinetochore (α-Ndc80), centrosome inner-core (α-HA marking Cep250L1-HA) and outer-core (α-Centrin) markers. Boxed areas magnified on the right.

## Discussion

In mammalian cells, duplicated centrosomes are tethered together by a fiber complex composed of C-NAP1/Cep250 ^21^, rootletin ^22^ and LRRC45 ^23^. Phosphorylation of C-NAP1/Cep250 by NIMA kinase Nek2 promotes degradation of C-NAP1/Cep250 and results in splitting of the linked centrosomes ^21^. Reciprocal BLAST analysis suggests that C-NAP1/Cep250 and rootletin orthologs are absent from the *Toxoplasma* genome ^6^, however, the regulatory machinery might be conserved.

We showed here that the functional ortholog of human Nek2 in *Toxoplasma*, TgNek1, is essential for outer-core splitting (Fig. 7) but not the inner-core. This observation implies that substrate of TgNek1 only tethers the outer-core and the mechanism that physically separates the inner- and outer-core is different. Both TgNek1 and TgCep250 are closely associated with the centrosome outer-core, however, their timing is different. TgNek1 is only transiently recruited to the centrosome in G1/S phase transition and disappears after cell cycle progresses into mitosis ^14^, which is just when TgCep250 shows up in the outer-core. Thus, although we cannot exclude the possibility of TgCep250 is a TgNek1 substrate, the proteolytic event of TgCep250 that leads to both inner- and outer-core localization is likely independent from TgNek1 activity since TgNek1 is already absent from the centrosome during cytokinesis. Phenotypic analysis of the TgCep250-cKD and TgNek1-cKD phenotypes also showed distinct cell cycle defects indicating the roles of TgCep250 and TgNek1 are functionally distinct. It is worth noticing that TgCep250 is phosphorylated at S252 in whole organism phopho-proteome data ^24^. However, the function of this TgCep250 phosphorylation and the kinase that phosphorylates it remains to be determined.

In mammalian cells, the differential decoration of centriole appendages defines the difference in age and ability to nucleate microtubules between the centriole pairs ^17^. *Toxoplasma*’s annotated genome does not contain orthologs of mammalian centriole appendage genes and no clear appendage structure was observed by transmission election microscopy. However, the observation that the detachment of outer-core and inner-core occurs mostly in one pair of centrosomes upon TgCep250 depletion provides the first evidence that daughter and mother centrosomes can be differentiated in apicomplexan parasites.

Overduplication of outer-cores has been reported in parasites expressing a temperature sensitive allele of PCM component MAPK-L1 ^7^, whereas outer-core fragmentation was observed in TgCep530 depleted parasites ^8^. In TgCep250 depleted cells we observed multiple outer-cores, which could either be the result of overduplication or fragmentation (Fig. 4C). The TgCep250 phenotypes differs from the MAPK-L1 defective cells as an accumulation of inner- and outer-cores aligned in a “dumbbell form” was not observed. On the other hand, the TgCep250 phenotype is also dramatically different from the multiple outer-cores of various intensity observed in TgCep530 deficient parasites. The multiple outer-cores observed upon TgCep250 depletion appear clearly organized and are of equal intensity, which is more consistent with an over duplication than with fragmentation. In addition, we observed normal initiation of daughter buds in absence of TgCep250 (Fig. 4D), suggestive of fully functional outer-cores.

The above observation provides further support for the hypothesis of independent function and regulation of inner- and outer-cores ^7^. In addition, the formation of spindle microtubules on disconnected inner-cores (Fig. 5A) is also in line with the independent functionality of the bifunctional centrosome cores. We further tested this model in an IFA assay combining SFA, Centrin, TgCep250L1 and TgNdc80 (Fig. 8), as the integrity of each of these detected structures is essential and critical for the endowment of each daughter cell with a single copy of genetic material ^4^, ^18^. Indeed, we observed a hierarchical organization of these structures required in wild type parasites (Fig. 8, upper panels). However, in TgCep250-cKD cells, the connection between the inner-(green) and outer-core (blue) is lost while association between the SFA and outer-core as well as between the kinetochore and the inner-core remain intact (Fig. 8, bottom panels).

**Figure 8.**
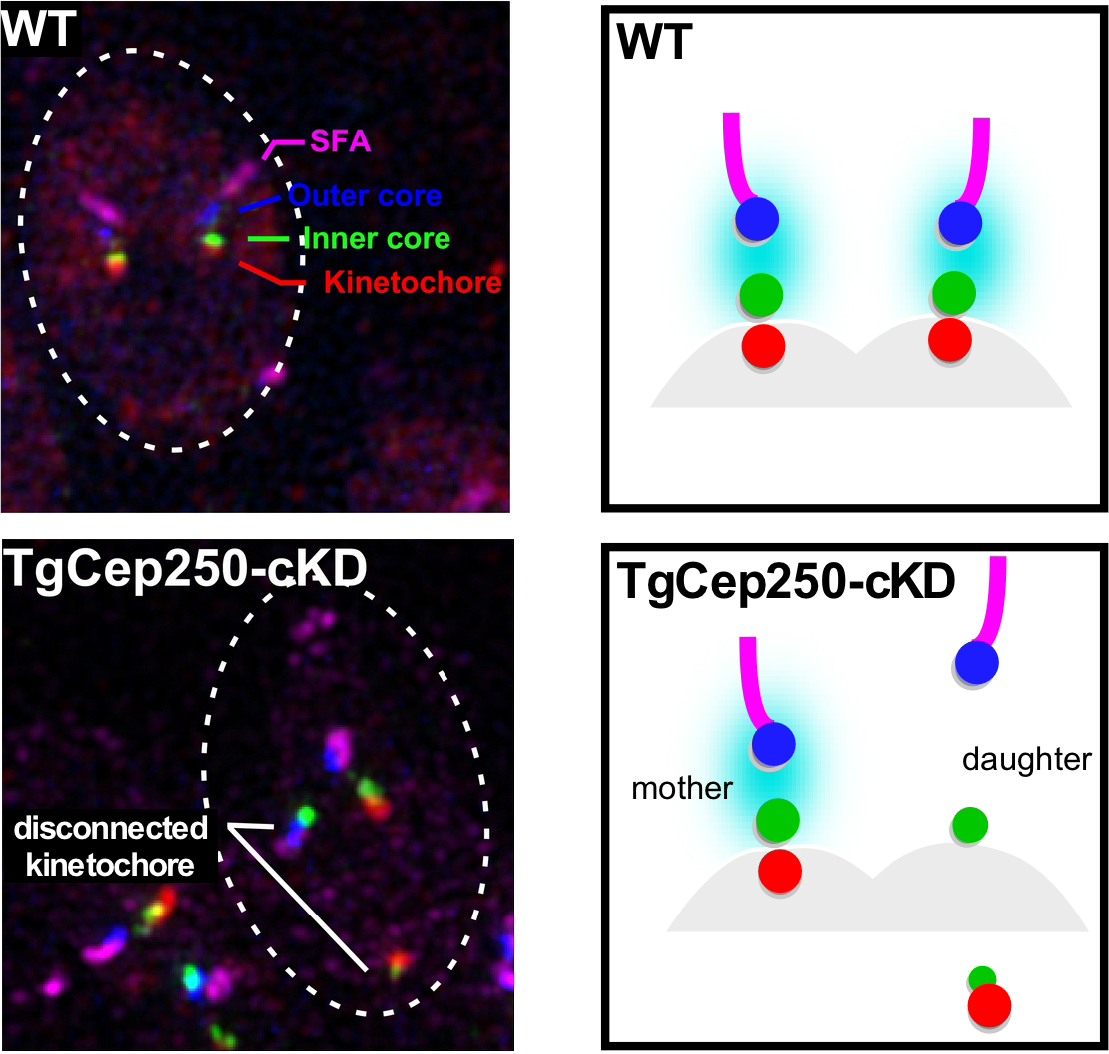
Hierarchical organization of the nuclear and cell division roles of the *Toxoplasma* centrosome inner- and outer-cores. Left panels: super-resolution (SIM) IFA of TgCep250-cKD co-expressing TgCep250L1-HA stained with α-HA (green: inner-core), α-Centrin (blue: outer-core), α-TgSFA (magenta: daughter tether) and α-TgNdc80 (red: kinetochore). Dotted lines outline parasites. Right panels represent schematic representation of the data with corresponding colors. Cyan represents the connection between the inner- and outer-core that is lost in the daughter centrosome upon depletion of TgCep250.

Our previous work showed that the kinetochore is required for Centrin (outer-core) association with nucleus ^20^ by locking the MTs emanating from the centrosome on the nuclear envelope ^11^. Using the novel tools developed here, we now expanded these insights by showing that in TgNuf2-cKD cells the intact inner/outer-core pair dissociates in its entirety from the nucleus (Fig. 6). Thus, we can conclude that the kinetochore serves as a molecular anchor to lock the complete centrosome cores, and that spindle MTs anchoring to the nuclear periphery is not required to maintain the connection between the inner- and outer-core.

Taken together, our data support the notion of functional independency of centrosomal cores as previous hypothesized. Furthermore, the observed differential states of mother and daughter centrosomes provide support for the non-geometric daughter expansion numbers observed in the *Plasmodium* spp. red blood cell cycle: it was proposed that the mother centrosome could be sooner ready for another round of duplication, associated with S-phase and mitosis, than the daughter centrosome, which still had to mature ^25^.

TgCep250 is essential for structurally stability of the connection between the centrosome cores and two different forms of Cep250 are present in tachyzoites: a full-length form localizing to the outer-core during the transition between mitosis/cytokinesis and a proteolytically processed form localizing to the inner-core throughout the tachyzoite cell cycle. TgNek1 is unlikely the kinase triggering proteolytic cleavage. However, the tethering function of TgCep250 appears to be maintained between human and *Toxoplasma* orthologs, but the structures connected are tailored to each organism. The phosphorylation controls and the proteolytic mechanisms remain unknown in *Toxoplasma* but provide an exciting area for future research.

## Material and Methods

### Parasite strains

*Toxoplasma gondii* tachyzoites of RHΔHX ^26^, RHΔKu80ΔHX ^16^, TATiΔKu80 ^27^ and their transgenic derivatives were grown in human foreskin fibroblast (HFF) as described ^28^. 1 μM pyrimethamine, 20 μM of chloramphenicol, or 25 μg/ml mycophenolic acid in combination with 50 μg/ml xanthine were used to select stable transgenic parasite lines and cloned by serial dilution. 1.0 μg/ml of anhydrotetracycline was used to repress specific gene expression of conditional knockdown lines. Cep250Like-HA fosmid was kindly provided by Dr. Michael White, University of South Florida ^29^. The *tub-EB1-YFP/sagCAT* plasmid has been described before ^11^.

### Plasmid constructs

All primers used are listed in Table S1. Endogenous tagging parasite strains were generated as described ^27^. Briefly, 1 kb and 1.9 kb region upstream of the stop codon of the *TgCep250* gene were PCR amplified using RH genomic DNA as template and were LIC-cloned into p*YFP-LIC-DHFR* (kindly provided by Dr. Vern Carruthers, University of Michigan). *Nar*I was used to linearize p*Cep250-YFP-LIC-DHFR* for site-specific homologous recombination.

Tetracycline regulatable parasite strains were generated by single-crossover homologous recombination using a plasmid kindly provided by Dr. Wassim Daher ^30^. The plasmid was re-designed by adding a single Myc-tag between the minimum promoter and the N-terminal genomic region of the target genes. All the inserts were amplified by PCR and digested with enzyme pairs *Bgl*II and *Not*I. To construct Myc-Cep250-cKD line, the 5’ end of the TgCep250 was amplified, digested with *Bgl*II and *Not*I enzyme pairs, and ligated into the DHFR-TetO7sag4-Myc vector. The resulting construct was linearized using NcoI, electroporated into the TATiΔKu80 line by electroporation, and selected for pyrimethamine resistance. The Myc-Nek1-cKD line was generated following the same strategy.

The Myc-Cep250-3xTy-cKD line was generated by site-specific insertion of a PCR product carrying the tag and HXGPRT selectable marker using CRISPR/Cas9. The pU6 plasmid expressing the guide RNA and Cas9 nuclease was kindly provided by Dr. Lourido from the Whitehead Institute ^31^. The CRISPR construct targeted the 3’UTR of TgCep250 near the stop codon. A primer pair was designed to amplify the 3xTy tag and the selectable marker (HXGPRT) flanking with 40 bp of homologous region prior to the stop codon and after the protospacer-adjacent motif (PAM) site. Amplicon generated from the primer pair was gel purified and co-transfected with the protospacer and Cas9 encoding plasmid.

### Immunofluorescence assay and microscopy

Parasites were seeded overnight (~16-18 h) and fixed with methanol as described ^9^. The following antibodies were used in this study: Myc (MAb 9E10, mouse, 1:50; Santa Cruz); HA (3F10, rat; 1:3000; Roche); IMC3 (rat, 1:2000 ^32^); TgNuf2 and TgNdc80 (guinea pig, 1:2000 ^20^); SFA (rabbit, 1:1000 ^13^ kindly provided by Dr. Boris Striepen, University of Georgia); TgNek1 (1:1000, ^14^), HsCentrin (rabbit, 1:1000; kindly provided by Dr. Iain Cheeseman, Whitehead Institute); Ty (mouse, 1:1000; kindly provided by Dr. Lourido). Alexa Fluor A488, A568, A594, A633, and A647 secondary antibodies were used. 4’,6’-diamidino-2-phenylindole (DAPI) was used to stain nuclear material. Images were acquired using a Zeiss Axiovert 200M wide-field fluorescent microscope equipped with DAPI, FITC, YFP and TRITC filter sets, and a Plan-Fluar 100×/1.45 NA oil objective and a Hamamatsu Orca-Flash 4.0LT camera. For SIM super-resolution microscopy a Zeiss Elyra S.1 microscope equipped with a Plan-Apochromat 63×/1.40 oil objective and PCO-Tech Inc. pco.edge 4.2 sCMOS camera in the Boston College Imaging Core was used in consultation with Bret Judson. Images were acquired and processed in Zeiss ZEN 2.3 software using standard mode.

### Western blot analysis

Freshy lysed parasites were collected after 3 μm filtration by centrifugation and washed twice in 1× PBS. Parasite pellets were lysed by resuspension in 50 mM Tris-HCl (pH7.8), 150 mM NaCl, 1% SDS containing 1× protease inhibitor cocktail (Sigma-Aldrich) and heating at 95°C for 10 min. Parasite lysates were analyzed by SDS-PAGE on a 3-8% Tris-acetate gradient gel. Prior to PVDF membrane transfer the gel was subjected to 100 mM acetic acid for 5 hr to break up the large TgCep250 protein. Transfer buffer contained 25 mM Tris pH 8.3, 195 mM glycine, 0.025% (w/v) SDS and 15% methanol. Blots were hybridized with α-Myc-HRP, α-Ty or α-α-tubulin (MAb 12G10; developmental studies hybridoma bank).

## Acknowledgements

We thank William Jacobus for technical assistance, Drs. Carruthers, Cheeseman, Daher, Lourido, Striepen, Suvorova, and White and for sharing reagents. We thank Bret Judson and the Boston College Imaging Core for infrastructure and support and Drs. White and Suvorova for fruitful suggestions and discussion.

This study was supported by National Science Foundation Major Instrumentation Grant 1626072 and through support from National Institute of Health grants AI110690, AI110638, and AI128136.

